# Contezolid can replace linezolid in a novel combination with bedaquiline and pretomanid in a murine model of tuberculosis

**DOI:** 10.1101/2023.06.18.545492

**Authors:** Deepak Almeida, Si-Yang Li, Jin Lee, Barry Hafkin, Khisimuzi Mdluli, Nader Fotouhi, Eric L. Nuermberger

**Affiliations:** Department of Medicine, Johns Hopkins University Center for Tuberculosis Research, Baltimore, Maryland, USA; MicuRx Pharmaceuticals, Foster City, California, USA; Global Alliance for Tuberculosis Drug Development, New York, New York, USA

## Abstract

Contezolid is a new oxazolidinone with *in vitro* and *in vivo* activity against *Mycobacterium tuberculosis* comparable to that of linezolid. Pre-clinical and clinical safety studies suggest it may be less toxic than linezolid, making contezolid a potential candidate to replace linezolid in treatment of drug-resistant tuberculosis. We evaluated the dose-ranging activity of contezolid, alone and in combination with bedaquiline and pretomanid, and compared it with linezolid at similar doses, in an established BALB/c mouse model of tuberculosis. Contezolid had an MIC of 1 µg/ml, similar to linezolid, and exhibited similar bactericidal activity in mice. Contezolid-resistant mutants selected *in vitro* had 10-fold increases in contezolid MIC and harbored mutations in the *mce3R* gene. These mutants did not display cross-resistance to linezolid. Our results indicate that contezolid has potential to replace linezolid in regimens containing bedaquiline and pretomanid and likely other regimens.

## Introduction

Despite recent advances in the development of new drugs against tuberculosis (TB), major challenges exist for the treatment of rifampin-resistant (RR-) TB. Linezolid (LZD), the first oxazolidinone antibacterial drug to reach the market has been repurposed to treat RR-TB and is now designated as a preferred drug for this indication by the WHO (1). LZD exhibited additive bactericidal and sterilizing activity when combined with bedaquiline (BDQ) and pretomanid (PMD) in mouse models of TB (2, 3). The resulting 3-drug “BPaL” regimen later proved to be an effective 6-month, oral regimen for extensively drug-resistant and refractory multidrug-resistant TB and received FDA approval for this indication. Despite the efficacy of the BDQ-PMD-LZD regimen, the dose- and duration-dependent toxicity of LZD compromises its utility (4-6). Replacing LZD with a newer oxazolidinones with similar activity and superior safety could reduce the burden of safety monitoring for treatment programs and expand the regimen’s spectrum of use to include drug-susceptible TB.

Contezolid (CZD), previously called MRX-I, is a newer oxazolidinone with potent activity against methicillin-, penicillin-, and vancomycin-resistant bacteria similar or superior to that of LZD (7). It was recently approved by the National Medical Products Administration in China for the treatment of complicated skin and soft tissue infections; and has already been used off-label in TB treatment (8). Although the MIC range of CZD against *Mycobacterium tuberculosis* (0.25-1.0 µg/ml) is similar to that of LZD (0.25-1.0 µg/ml), it may have a superior safety profile (7, 9, 10).

Here, we tested whether CZD could replace LZD in the BDQ-PMD-LZD regimen without loss of efficacy in a well-established mouse model of TB. CZD was used in a dose-ranging fashion, alone and in combination with BDQ-PMD, to evaluate its bactericidal activity relative to that of LZD. We also selected and characterized CZD-resistant mutants *in vitro* and evaluated them for cross-resistance with LZD.

## Results

### MIC determination

MICs of CZD and LZD against the parent *M. tuberculosis* H37Rv strain were determined using the broth macrodilution method and found to be 0.5-1 and 0.5-1 µg/ml, respectively.

### Efficacy in mouse model of TB

The dose-ranging activity of CZD and LZD, both alone and in combination with BDQ and PMD, was evaluated in a subacute BALB/c mouse infection model. Results of lung colony-forming unit (CFU) counts are shown in Table 1. After four weeks of treatment, dose-dependent activity was observed for both oxazolidinones. Mice in the LZD 100 mg/kg and CZD 100 and 200 mg/kg monotherapy groups had significantly lower mean CFU counts compared to pre-treatment (D0) controls (P < 0.05 for CZD at 100 mg/kg; P ≤ 0.0001 for others), while LZD and CZD at 50 mg/kg were largely bacteriostatic. Addition of each oxazolidinone increased the bactericidal activity of BDQ-PMD, although only LZD 100 mg/kg and CZD at 100 or 200 mg/kg increased the activity of the BDQ-PMD combination in a statistically significant fashion (P < 0.05 for CZD at 100 mg/kg; P < 0.001 for others). BDQ-PMD plus CZD at 200 mg/kg produced the lowest mean CFU count, but was not statistically superior to BDQ-PMD-LZD_100_.

**Table 1:** Mean lung CFU counts observed in the mouse efficacy experiment

| Regimen* | Mean (± SD) $\log_{10}$ CFU/lung | | |
| --- | --- | --- | --- |
|  | Day -13 | Day 0 | Week 4 |
| Untreated | 3.59 ± 0.07 | 6.21 ± 0.18 | |
| LZD<sub>50</sub> | | | 6.09 ± 0.23 |
| LZD<sub>100</sub> | | | 5.37 ± 0.16 |
| CZD<sub>50</sub> | | | 6.15 ± 0.22 |
| CZD<sub>100</sub> | | | 5.78 ± 0.31 |
| CZD<sub>200</sub> | | | 5.36 ± 0.25 |
| BDQ-PMD | | | 1.39 ± 0.12 |
| BDQ-PMD-LZD<sub>50</sub> | | | 0.96 ± 0.16 |
| BDQ-PMD-LZD<sub>100</sub> | | | 0.52 ± 0.20 |
| BDQ-PMD-CZD<sub>50</sub> | | | 1.16 ± 0.44 |
| BDQ-PMD-CZD<sub>100</sub> | | | 0.81 ± 0.32 |
| BDQ-PMD-CZD<sub>200</sub> | | | 0.42 ± 0.32 |

\*Subscripts indicate the LZD or CZD dose in mg/kg body weight

### Selection and characterization of drug-resistant mutants

To evaluate the potential for cross-resistance between CZD and LZD, mutants resistant to both drugs were selected *in vitro* and whole genome sequencing was performed to identify mutations. The calculated frequency of resistance was approximately 2×10^−5^ and 4×10^−6^ at 3 and 10 µg/ml of CZD, respectively, compared to 3×10^−8^ for LZD. Each of the resistant colonies sequenced from the CZD-containing plates (1 from each CZD concentration) had acquired a stop codon mutation in the *mce3R* gene, affecting either the Tyr185 or the Tyr118 residue. No other new mutations were found to be in common between the resistant isolates. These mutants exhibited elevated CZD MICs of 32-64 µg/ml, but retained full susceptibility to LZD (Table 2). In contrast, mutants selected on LZD-containing plates harbored either *rplC* (nt460 T→C; C154R) or *rrl* (nt2814 G→T) mutations. Both of these LZD-resistant mutants showed cross-resistance to CZD.

**Table 2:** Linezolid and contezolid MICs against the *M. tuberculosis* H37Rv parent strain and resistant mutants selected *in vitro*

| Strain | LZD MIC (µg/ml)* | CZD MIC (µg/ml) |
| --- | --- | --- |
| 1) H37Rv parent | 0.5-1 | 0.5-1 |
| 2) *mce3R* mutant selected on<br>CZD 3 µg/ml | 1 | 32 |
| 3) *mce3R* mutant selected on<br>CZD 10 µg/ml | 1 | 64 |
| 4) *rplC* C154R mutant selected<br>on LZD | 16-32 | 16 |
| 5) *rrl* nt2814 G→T mutant<br>selected on LZD | 16-32 | 32 |

\*LZD MICs against strains 1, 4 and 5 and CZD MIC against strain 1 were tested twice and the MIC range is shown.

## Discussion

Short-course oral regimens based on the BDQ-PMD-LZD combination have demonstrated high rates of successful clinical outcomes in the treatment of RR-TB, but dose- and duration-dependent toxicity of LZD remains a challenge (4-6). There is great interest in identifying a safer and equally efficacious oxazolidinone to replace LZD. CZD is an orally bioavailable oxazolidinone recently approved in China for treatment of complicated skin and soft tissue infections. In a phase 1 trial, no hematological adverse events were observed with CZD administered at 800 mg twice daily for 28 days (n=10), whereas 2 out of 10 subjects receiving LZD 600 mg twice daily developed hematological adverse events (7, 10). When the same dosing regimens were compared over 7-14 days in a phase 3 trial, CZD was associated with significantly fewer hematological adverse events than LZD (11). An ongoing phase 3 trial will compare the safety and efficacy of CZD and LZD administered for 14-28 days in treatment of diabetic foot infection (ClinicalTrials.gov Identifier: NCT05369052). Although LZD is commonly used at 600 mg once daily for treatment of TB, adverse effects still require vigilant safety monitoring and frequently limit the dose and duration of LZD use. Based on the clinical studies performed to date, CZD could represent a better oxazolidinone for TB treatment if it is at least as effective as LZD at exposures that are safer.

As CZD demonstrated potency comparable to LZD against *M. tuberculosis in vitro* and *in vivo*, including similar efficacy to LZD when tested in an acute mouse infection model (9), we investigated the replacement of LZD with CZD in the short-course oral regimen of BDQ, PMD and LZD. All tested CZD doses showed equivalent efficacy when compared to the same dose of LZD. Available data suggest that CZD and LZD produce similar exposures in mice at the same dose, as measured by the area under the plasma concentration-time curve (AUC) after oral administration. For example, a single oral dose of CZD 90 mg/kg in mice resulted in an average AUC_0-24h_ of 200.8 µg-h/ml (12), comparable to an average AUC_0-24h_ of 243.5 µg-h/ml after a single oral dose of LZD 100 mg/kg (2). Assuming similar exposures of CZD and LZD were achieved at the same doses in our study, our results indicate that CZD may be a suitable candidate to replace LZD in RR-TB treatment, including regimens with BDQ and PMD, if CZD is safer in humans at exposures as high or higher than efficacious LZD exposures.

Studies against *S. aureus* showed extremely low frequencies of spontaneous resistance to CZD and LZD between 10^−12^ and 10^−13^ (13), with resistance caused by mutations in the 23S rRNA and *rplC* genes. Mutations in these genes are known to confer LZD resistance in *M. tuberculosis* at a frequency of 1×10^−8^ (14). The spontaneous frequency of resistance to CZD in *M. tuberculosis* was unexpectedly higher than 10^−6^, or roughly 100 times higher than we observed for LZD. Whole genome sequence analysis of CZD-selected mutants indicated that the higher frequency of spontaneous resistance was likely driven by mutations in *mce3R*. These results now independently confirm the recently published results from Pi et al (15), which demonstrated that a variety of loss-of-function mutations in *mce3R* confer resistance to CZD, likely through de-repression of a drug-modifying flavin-dependent mono-oxygenase encoded by *Rv1936*. Consistent with the observed difference in the frequency of spontaneous resistance between CZD and LZD, the CZD-resistant *mce3R* mutant was not cross-resistant to LZD. This finding, also noted by Pi et al (15), suggests the intriguing possibility that, should CZD prove safe and effective enough to replace LZD as the oxazolidinone of choice in TB regimens, emergence of resistance to CZD via the more frequent *mce3R* mutation would not obviate the future utility of LZD in a re-treatment regimen. Nevertheless, the higher spontaneous frequency of resistance observed for CZD compared to LZD would theoretically place a greater demand on companion drugs to prevent the selection of CZD-resistant mutants during combination therapy.

In conclusion, CZD is novel oxazolidinone that is as efficacious as LZD at similar doses in murine models of TB and could prove to be a safer alternative to LZD for TB treatment.

## Materials and Methods

### Mycobacterial strains

*M. tuberculosis* H37Rv (ATCC 27294) was passaged in mice, sub-cultured in Middlebrook 7H9 broth supplemented with 10% (v/v) oleic acid-albumin-dextrose-catalase (OADC) enrichment (Becton-Dickinson) and 0.05% (v/v) Tween 80 (Fisher Scientific), and used for aerosol infection when the optical density at 600 nm was approximately 1.0.

### Antimicrobials

CZD and BDQ were provided by MicuRx Pharmaceuticals and Janssen Pharmaceuticals, respectively. LZD and PMD were provided by the Global Alliance for Tuberculosis Drug Development (New York, NY). Dosing formulations were prepared and maintained as previously described for PMD (16, 17), BDQ and LZD (18). CZD was prepared in 0.5% carboxymethylcellulose sodium (CMC-Na). All drugs were administered once daily by gavage, 5 days per week; when given in combination, BDQ and PMD were given one after another and LZD or CZD were given 4 hours later.

### MIC determination

*M. tuberculosis* H37Rv was grown in Middlebrook 7H9 broth supplemented with 10% OADC (BD), 0.5% glycerol and 0.05% Tween. CZD and LZD stock solutions were prepared in 100% DMSO. Serial 2-fold dilutions of each stock solution were made with the same 7H9 media but without Tween 80 to yield final assay concentrations ranging from 0.06 to 64 µg/ml. Drug-free tubes served as growth controls. Each polystyrene tube containing 2.4 ml of 2-fold diluted drug solution was inoculated with 100 µl of *M. tuberculosis* culture adjusted to 0.1 OD at 600 nm to give a final bacterial concentration of approximately 10^5^ CFU-ml^-1^. Initial assessment was made after 1 week of incubation, followed by a final assessment at 2 weeks. The MIC was defined as the lowest concentration that inhibited visible bacterial growth after 14 days of incubation at 37°C.

### Mouse infection model

Female BALB/c mice (Charles River, Wilmington, MA) aged 4 to 6 weeks were infected with the goal of implanting 3.5-4 log_10_ CFU of *M. tuberculosis* H37Rv by the aerosol route using an inhalation exposure system (Glas-col Inc., Terre Haute, IN). Two mice were sacrificed one day after infection (D-13) to determine the number of CFU implanted in the lungs and 5 more were sacrificed on the day of treatment initiation (D0) to determine the pre-treatment CFU count.

### Drug treatment

Treatment started 2 weeks after infection (D0) with 5 mice randomized to each treatment group. Test mice received CZD at 50, 100, or 200 mg/kg, alone or in combination with BDQ at 25 mg/kg and PMD at 100 mg/kg. Control mice went untreated or received LZD at 50 or 100 mg/kg, alone or in combination with BDQ-PMD, or BDQ-PMD alone. All treated mice were sacrificed after 4 weeks of treatment to determine the efficacy of treatment. Lung CFU counts were determined by plating 0.5 ml aliquots of serially diluted lung homogenates on Middlebrook 7H11 selective plates containing 0.4% activated charcoal to minimize the effects of drug carryover (18). All procedures involving animals were approved by the Animal Care and Use Committee of Johns Hopkins University.

### Statistical analysis

Mean lung CFU counts were compared using one-way ANOVA with Dunnett’s post-test (GraphPad Prism).

### Selection and characterization of drug-resistant mutants

A total inoculum of between 10^8^ and 10^9^ CFU was plated on 7H11 agar plates containing CZD at 3 or 10 µg/ml or LZD at 1.5 µg/ml. After 4 weeks of incubation, individual colonies growing on drug–containing plates were selected for confirmatory MIC testing. Genomic DNA from colonies selected on CZD was subjected to whole genome sequencing at the Johns Hopkins University Next Generation Sequencing Center. Samples were prepared using the TruSeq DNA kit. Sequencing was performed on an Illumina HS2500. Alignments were done by bwa0.7.7 and data aligned to H37Rv genome NC_000962.3. Samtools1.2.1 (mpileup) was used to call variants. snpEFF4.1 was used to annotate the variant calls. Genomic DNA from colonies selected on LZD-containing plates was subjected to PCR to amplify the *rplC* and *rrl* genes known to confer LZD resistance in *M. tuberculosis*, using previously described methods (19).

## Acknowledgements

Financial support was provided by the Global Alliance for TB Drug Development. CZD was provided by MicuRx Pharmaceuticals, Inc.

## References

1. WHO. 2019. WHO consolidated guidelines on drug-resistant tuberculosis treatment. World Health Organization, Geneva. http://www.who.int/tb/publications/2019/consolidated-guidelines-drug-resistant-TB-treatment/en/.

2. Tasneen, R., F. Betoudji, S. Tyagi, S. Li, K. Williams, P. J. Converse, V. Dartois, T. Yang, C. M. Mendel, K. E. Mdluli, and E. L. Nuermberger. 2016. Contribution of Oxazolidinones to the Efficacy of Novel Regimens Containing Bedaquiline and Pretomanid in a Mouse Model of Tuberculosis. Antimicrob. Agents Chemother. 60:270–277. doi: 10.1128/AAC.01691-15. http://aac.asm.org/content/60/1/270.abstract.

3. Xu, J., S. Y. Li, D. V. Almeida, R. Tasneen, K. Barnes-Boyle, P. J. Converse, A. M. Upton, K. Mdluli, N. Fotouhi, and E. L. Nuermberger. 2019. Contribution of pretomanid to novel regimens containing bedaquiline with either linezolid or moxifloxacin and pyrazinamide in murine models of tuberculosis. Antimicrob. Agents Chemother.. doi: AAC.00021-19 [pii].

4. Sotgiu, G., R. Centis, L. D’Ambrosio, J. W. Alffenaar, H. A. Anger, J. A. Caminero, P. Castiglia, S. De Lorenzo, G. Ferrara, W. J. Koh, G. F. Schecter, T. S. Shim, R. Singla, A. Skrahina, A. Spanevello, Z. F. Udwadia, M. Villar, E. Zampogna, J. P. Zellweger, A. Zumla, and G. B. Migliori. 2012. Efficacy, safety and tolerability of linezolid containing regimens in treating MDR-TB and XDR-TB: systematic review and meta-analysis. Eur. Respir. J. 40:1430–1442. doi: 10.1183/09031936.00022912 [doi].

5. Conradie, F., T. R. Bagdasaryan, S. Borisov, P. Howell, L. Mikiashvili, N. Ngubane, A. Samoilova, S. Skornykova, E. Tudor, E. Variava, P. Yablonskiy, D. Everitt, G. H. Wills, E. Sun, M. Olugbosi, E. Egizi, M. Li, A. Holsta, J. Timm, A. Bateson, A. M. Crook, S. M. Fabiane, R. Hunt, T. D. McHugh, C. D. Tweed, S. Foraida, C. M. Mendel, M. Spigelman, and ZeNix Trial Team. 2022. Bedaquiline-Pretomanid-Linezolid Regimens for Drug-Resistant Tuberculosis. N. Engl. J. Med. 387:810–823. doi: 10.1056/NEJMoa2119430.

6. Conradie, F., A. H. Diacon, N. Ngubane, P. Howell, D. Everitt, A. M. Crook, C. M. Mendel, E. Egizi, J. Moreira, J. Timm, T. D. McHugh, G. H. Wills, A. Bateson, R. Hunt, C. Van Niekerk, M. Li, M. Olugbosi, M. Spigelman, and Nix-TB Trial Team. 2020. Treatment of Highly Drug-Resistant Pulmonary Tuberculosis. N. Engl. J. Med. 382:893–902. doi: 10.1056/NEJMoa1901814.

7. Gordeev, M. F., and Z. Y. Yuan. 2014. New potent antibacterial oxazolidinone (MRX-I) with an improved class safety profile. J. Med. Chem. 57:4487–4497. doi: 10.1021/jm401931e [doi].

8. Kang, Y., C. Ge, H. Zhang, S. Liu, H. Guo, and J. Cui. 2022. Compassionate Use of Contezolid for the Treatment of Tuberculous Pleurisy in a Patient with a Leadless Pacemaker. Infect. Drug Resist. 15:4467–4470. doi: 10.2147/IDR.S373082.

9. Shoen, C., M. DeStefano, B. Hafkin, and M. Cynamon. 2018. In Vitro and In Vivo Activities of Contezolid (MRX-I) against Mycobacterium tuberculosis. Antimicrob. Agents Chemother. 62:10.1128/AAC.00493-18. Print 2018 Aug. doi: e00493-18 [pii].

10. Eckburg, P. B., Y. Ge, and B. Hafkin. 2017. Single- and Multiple-Dose Study To Determine the Safety, Tolerability, Pharmacokinetics, and Food Effect of Oral MRX-I versus Linezolid in Healthy Adult Subjects. Antimicrob. Agents Chemother. 61:10.1128/AAC.02181-16. Print 2017 Apr. doi: e02181-16 [pii].

11. Zhao, X., H. Huang, H. Yuan, Z. Yuan, and Y. Zhang. 2022. A Phase III multicentre, randomized, double-blind trial to evaluate the efficacy and safety of oral contezolid versus linezolid in adults with complicated skin and soft tissue infections. J. Antimicrob. Chemother. 77:1762–1769. doi: 10.1093/jac/dkac073.

12. Chen, X. Y., Gao, Z. W., Yuan, H., Yuan, Z. Y., Zhong D. F. 2012. Abstr 52nd Intersci Conf Antimicrob Agents Chemother, abstr F-1500.

13. Huang, Y., Y. Xu, S. Liu, H. Wang, X. Xu, Q. Guo, B. Wu, M. F. Gordeev, W. Wang, Z. Yuan, and M. Wang. 2014. Selection and characterisation of Staphylococcus aureus mutants with reduced susceptibility to the investigational oxazolidinone MRX-I. Int. J. Antimicrob. Agents. 43:418–422. doi: 10.1016/j.ijantimicag.2014.02.008 [doi].

14. Pi, R., Q. Liu, Q. Jiang, and Q. Gao. 2019. Characterization of linezolid-resistance-associated mutations in Mycobacterium tuberculosis through WGS. J Antimicrob Chemother. 74:1795–1798. doi: 10.1093/jac/dkz150. https://academic.oup.com/jac/article/74/7/1795/5477395.

15. Pi, R., X. Chen, J. Meng, Q. Liu, Y. Chen, C. Bei, C. Wang, and Q. Gao. 2022. Drug Degradation Caused by mce3R Mutations Confers Contezolid (MRX-I) Resistance in Mycobacterium tuberculosis. Antimicrob. Agents Chemother. 66:e0103422. doi: 10.1128/aac.01034-22.

16. Tyagi, S., E. Nuermberger, T. Yoshimatsu, K. Williams, I. Rosenthal, N. Lounis, W. Bishai, and J. Grosset. 2005. Bactericidal activity of the nitroimidazopyran PA-824 in a murine model of tuberculosis. Antimicrob. Agents Chemother. 49:2289–2293. doi: 49/6/2289 [pii].

17. Nuermberger, E., I. Rosenthal, S. Tyagi, K. N. Williams, D. Almeida, C. A. Peloquin, W. R. Bishai, and J. H. Grosset. 2006. Combination chemotherapy with the nitroimidazopyran PA-824 and first-line drugs in a murine model of tuberculosis. Antimicrob. Agents Chemother. 50:2621–2625. doi: 50/8/2621 [pii].

18. Tasneen, R., S. Li, C. A. Peloquin, D. Taylor, K. N. Williams, K. Andries, K. E. Mdluli, and E. L. Nuermberger. 2011. Sterilizing Activity of Novel TMC207- and PA-824-Containing Regimens in a Murine Model of Tuberculosis. Antimicrobial Agents and Chemotherapy. 55:5485–5492. doi: 10.1128/AAC.05293-11. http://aac.asm.org/content/55/12/5485.abstract.

19. Zhang, S., J. Chen, P. Cui, W. Shi, X. Shi, H. Niu, D. Chan, W. W. Yew, W. Zhang, and Y. Zhang. 2016. Mycobacterium tuberculosis Mutations Associated with Reduced Susceptibility to Linezolid. Antimicrob. Agents Chemother. 60:2542–2544. doi: 10.1128/AAC.02941-15.

